# Ribosome profiling uncovers selective mRNA translation associated with eIF2 phosphorylation in erythroid progenitors

**DOI:** 10.1101/216234

**Authors:** Nahuel A. Paolini, Kat S. Moore, Franca M. di Summa, Ivo F.A.C. Fokkema, Peter A.C. ‘t Hoen, Marieke von LIndern

**Affiliations:** Department of Hematopoiesis, Sanquin Research, and Landsteiner Laboratory AMC/UvA, 1066 CX Amsterdam, The Netherlands; Department of Human Genetics, Leiden University Medical Center, 2300 RC Leiden, The Netherlands

## Abstract

The regulation of translation initiation factor 2 (eIF2) is important for erythroid survival and differentiation. Lack of iron, a critical component of heme and hemoglobin, activates Heme Regulated Inhibitor (HRI). This results in phosphorylation of eIF2 and reduced eIF2 availability, which inhibits protein synthesis. Translation of specific transcripts such as *Atf4*, however, is enhanced. Upstream open reading frames (uORFs) are key to this regulation. The aim of this study is to investigate how eIF2 phosphorylation affects mRNA translation in erythroblasts. Ribosome profiling combined with RNA sequencing was used to determine translation initiation sites and ribosome density on individual transcripts. Treatment of erythroblasts with Tunicamycin (Tm) increased phosphorylation of eIF2 2-fold. At a false discovery rate of 1%, ribosome density was increased for 147 transcripts, among which transcriptional regulators such as *Atf4*, *Tis7/Ifrd1*, *Pnrc2*, *Gtf2h*, *Mbd3*, *JunB* and *Kmt2e*. Translation of 337 transcripts decreased more than average, among which *Dym* and *Csde1*. Ribosome profiling following Harringtonine treatment uncovered novel translation initiation sites and uORFs. Surprisingly, translated uORFs did not predict eIF2-dependent translation efficiency, but uORF identity differs. The regulation of transcription and translation factors in reponse to eIF2 phosphorylation may explain the large overall response to iron deficiency in erythroblasts.

- eif2 dependent translation in erythroblasts during proteotoxic stress determined by ribosome footprinting
- identification of transcription factors upregulated in response to eIF2 phosphorylation
- Advantages and disadvantages of translation initiation site determination using harringtonine
- distinct uORF pattern in transcripts with enhanced, or more than average reduced translation upon proteotoxic stress

## Introduction

Iron is an important element for life, but its strong reducing capacity is also very toxic and could create oxidative radicals in the cell [1], Therefore, the uptake of iron from the diet is limited, and circulating iron is always bound to carriers. Iron is a rate limiting factor for the production of hemoglobin during red blood cell development [2], Iron deficiency reduces heme availability, and risks the accumulation and aggregation of free α- and β-globin proteins that damage the cell [3], Therefore, it is important that globin protein synthesis is adjusted to iron availability. The Iron response element binding proteins Irp1 and Irp2 control mRNA stability and translation of transcripts encoding proteins involved in iron homeostasis such as the Transferrin receptor, Ferroportin, and Ferritin [4], Animal models for iron deficiency anemia, or iron depletion upon blood donation, indicate that not only differentiation, but also expansion of immature erythroblasts is impaired [5,6], The cellular mechanism responsible for impaired erythroid recovery upon iron deficiency, however, is poorly understood.

Reduced availability of heme also activates eIF2 associated kinase 1 (eIF2ak1, also known as HRI [Heme Regulated Inhibitor]) that phosphorylates translation initiator factor 2α (eIF2α) [7], GTP-bound eIF2 and methionine-loaded initiatior tRNA (tRNA_i_^met^) form the ternary complex (TC). The TC binds to the 40S small ribosomal subunit in the preinitiation scanning complex. The GTPase activity of eIF2 is activated when the scanning complex pauses at a translation start site, which results in release of methionine to the P-site of the ribosome, and dissociation of both tRNA_i_ and GDP-bound eIF2 from the scanning complex [8], The GDP-GTP exchange factor eIF2B reloads eIF2 with GTP, which enables eIF2 to bind tRNA_i_^met^ and to re-associate with a preinitiation scanning complex. Phosphorylation of the α-chain of eIF2 (eIF2α) on Ser51 by HRI prevents exchange of GDP for GTP and thereby recovery of the TC. As a result protein synthesis is inhibited to decrease globin production, which prevents damage from globin protein aggregates [9], Three additional kinases are able to phosphorylate eIF2α: the double-stranded RNA-dependent kinase (PKR, or Eif2ak2), the ER-stress activated kinase PERK (Eif2ak3), and GCN2 (general control nonderepressible 2, or Eif2ak4) that is activated by uncharged tRNA upon lack of amino acids [10].

Translational control by eIF2 is, at least in part, mediated through translation of upstream open reading frames (uORFs). Whereas general translation is repressed, translation of specific transcripts is increased upon eIF2 phosphorylation, as described for *Atf4*. A distance of ~90 nt between the first and second uORF allows for re-association in absence of eIF2 phosphorylation [11], Translation of the second uORF overlapping the start codon of the protein coding ORF inhibits Atf4 protein expression. Reduced availability of eIF2 decreases translation initiation at the second uORF (also referred to as leaky scanning), and increases translation of the *Atf4* protein coding ORF. The short distance between uORFs is crucial for eIF2-mediated control of translation [11,12], Phosphorylation of eIF2 also reduces the recognition of start codons in a suboptimal Kozak consensus context as is exemplified by the regulation of *Ddit3* (Death and differentiation induced transcript 3, also known as Chop). The inhibitory uORF of *Ddit3* is poorly translated upon eIF2 phosphorylation, which increases Ddit3 protein expression [13], Depending on the configuration of the 5’UTR, translation of specific transcripts can also be hypersensitive for eIF2 and cause a more than average repression of translation, as has been described for *Csde1* [14].

Whereas these examples demonstrate quantitative effects on protein synthesis, uORFs are also involved in qualitative changes in protein expression. A short distance between an uORF and the start codon of the protein coding ORF may result in partial availability of the protein initiating start codon. The presence of a downstream, in frame, start codon can subsequently result in expression of an N-terminally truncated short isoform. This leaky scanning controls for instance the balance between the long and short isoform of Tal1/Scl, an important transcription factor in erythropoiesis [15].

Heme-regulated phosphorylation of eIF2 and the subsequent regulation of mRNA translation, is important in the control of erythropoiesis. HRI-induced expression of Atf4 and its downstream target Ppp1r15a/Gadd34 constitutes an integrated stress response (ISR) that protects erythroid progenitors from oxidative stress during differentiation, and increases survival of erythroid cells when mice are fed a low iron diet [16], Atf4 null mice displayed severe fetal anemia [17], Modulation of the stress response is regulated by the dephosphorylation of eIF2 by Ppp1r15a and Ppp1r15b [18,19], Loss of Ppp1r15a results in enlarged spleens with increased numbers of immature erythroid cells and low hemoglobin content [20], Loss of Ppp1r15b increases the number of deformed erythroblasts and reduces the number of mature erythrocytes. The erythrocyte numbers were rescued when loss of Ppp1r15b was combined with the S51A knock-in mutation of eIF2, that abrogates eIF2 phosphorylation [21], These phenotypes indicate that eIF2 phosphorylation is important for control of both expansion and differentiation of erythroblasts.

The importance of translational control in erythropoiesis was demonstrated by polyribosome profiling, which allows for the identification of RNA populations in distinct fractions of a density gradient that separates subpolysomal RNA from low and high density polyribosomes [22,23], Ribosome footprinting or ribo-seq allows for deep sequencing of mRNA fragments protected by the ribosome (ribosome footprints, RFPs) [24,25], The RFPs are aligned to the genome, which maps the position of ribosomes at the nucleotide level and adds considerable detail to the analysis of mRNA translation. The aim of this study is to identify transcripts that are hypersensitive to eIF2 phosphorylation, and that encode proteins controlling expansion and differentiation of erythroblasts. We hypothesize that translation of uORFs renders transcripts sensitive to eIF2 phosphorylation because it controls re-association of the TC with the preinitiation scanning complex, which is required for translation of a subsequent ORF. We aim to identify cellular mechanisms regulated by eIF2 phosphorylation that are involved in erythroid homeostasis. We employed ribosome footprint analysis in combination with mRNA sequencing to identify both translation initiation sites (TIS) and the relative translation efficiency of transcripts. At a false discovery rate (FDR) of 1% we identified 147 transcripts subject to increased translation, and 337 transcripts subject to reduced translation upon eIF2 phosphorylation. Interestingly, the presence of translated uORFs was widespread, but did not predict sensitivity of the mRNA translation to eIF2 phosphorylation. Among the transcripts subject to eIF2-dependent translation were several transcription factors that may alter programming of erythropoiesis upon eIF2 phosphorylation.

## Materials and methods

### Cell culture

The erythroblast cell line 15.4 was derived from p53-deficient mouse fetal livers as previously described [26], and cultured in Stempro-34 SFM (Thermo Fisher), containing penicilin-streptavidin, L-glutamin, Erythropoietin (1U/ml), Stem Cell Factor (supernatant CHO cells) and 1μM Dexamethasone (Sigma) [27], For ER stress induction, cells were treated with 2.5μ/ml Tunicamycin (Tm) (Sigma) for 1.5h or left untreated. *SDS-PAGE*. Whole cell lysates were loaded on 10% polyacrylamide gels (Biorad). Western blots were performed as previously described [22], Antibodies used were eIF2 (Cell Signaling) and pSer51-eIF2 (Cell Signaling).

### Polysome profiling

10^7^ cells were lysed in polysome lysis buffer (110 mM potassium acetate, 20 mM magnesiumacetate, 10 mM HEPES, 100 mM potassium chloride, 10 mM magnesium chloride, 0.1% NP-40, 2 mM DTT, 40 U/mL RNase inhibitor [Thermo Fisher], 100 μg/ml cycloheximide [CHX] [Sigma] and 1X mini Protease Inhibitor Cocktail [Roche]) and loaded onto 17-50% sucrose gradients [28], The tubes were centrifuged at 40,000 rpm for 2 hours at 4°C in a SW41 rotor (Optima L100XP ultracentrifuge; Beckman Coulter). RNA was measured throughout the gradient with a BR-188 Density Gradient Fractionation System at OD_254_ (Brandel). Area under the curve was calculated with Fiji, statistical significance was calculated with a t-test. P-values < 0.01 were considered significant.

### Measurement of de novo protein synthesis

100,000 erythroblasts were seeded in methionine-free DMEM (Invitrogen) for 60 minutes to deplete intracellular methionine, followed by a 90 minutes exposure to Click-iT^®^ AHA (a methionine analogue) in absence or presence of 2.5μg/ml Tm treatment. Newly synthesised protein was measured using the Click-iT^®^ AHA Alexa Fluor^®^ 488 Protein Synthesis HCS Assay (Thermo Scientific) according to manufacturer’s instructions with some modifications. Briefly, using 2% paraformaldehyde for fixation and 1:1000 dilution of AHA. Fluorescence was measured by using an LSR-II flow cytometer and analyzed with FACSDiva software (BD Biosciences).

### Ribosome profiling and RNAseq

The ribosome profiling strategy was adapted from Ingolia et al. [30] and based on De Klerk et al. [31], with some modifications. After Tm treatment, the cells were treated with 100 μg/ml cycloheximide (CHX) for 5 min at 37°C or 2 μg/ml Harringtonine for 7 min followed by 2 min 100 μg/ml CHX at 37°C. Cells were lysed in polysome lysis buffer. Lysates were treated with 1500 units of RNAse-l (Ambion) to digest the polysomes into monosomes. The 80S monosome fraction was isolated by ultracentrifugation (Beckman) on sucrose gradients and RNA was isolated as described [31]. Ribosomal RNA (rRNA) was removed with Ribozero Gold rRNA Removal Kit (Illumina). In this study, the Nebnext small RNA Library Prep Set for Illumina (NEB) was used, according to manufacturer’s instructions and sequenced on a HiSeq Illumina. For RNAseq, mRNAs with a Poly-A tail were isolated, fragmented and sequenced on a Hiseq Illumina using the Truseq protocol.

### Data analysis

Adapters were trimmed with cutadapt [32], Reads were mapped to the transcriptome and unaligned reads to the genome with Spliced Transcripts Alignment to Reference (STAR) version 2.5.2b [33] with the following settings: --outFilterMultimapNmax 20 --outFilterMismatchNmax 1 –outSAMmultNmax 1. A GTF annotation file accessed from the UCSC genome browser on ll-Sept-2015 was passed to STAR to improve mapping accuracy. Translation efficiency was determined using the Bioconductor package edgeR (Empirical Analysis of Gene Expression Data in R) [34,35], edgeR utilizes a negative binomial distributed model for each gene and sample, scaled by library size and relative abundance per experimental group. An empirical Bayes procedure is applied to shrink dispersions towards a consensus value. Ribosome density was estimated via the application of a generalized linear model to determine the interaction between sequence assay (ribosome profiling versus RNA-seq) and condition (Tm-treated versus untreated) while also taking variation between different independent replicate experiments (performed on three different days) into account, using the formula expression level ~ replicate + condition*type + error. The application of an interaction term is a statistically formalized way of detecting which transcripts are translated with different efficiencies upon Tm treatment, as their level of active translation (ribosome profiling) will respond differently to Tm treatment than their total RNA levels (RNA-seq).

Prior to statistical analysis, ribosome footprint reads were separated based on their position in the 5’UTR, the protein coding ORF of the reference transcript 1 (CDS), or the 3’UTR. We did not correct for mapping a read to the first nucleotide of the protected fragment, which was position −13 compared to the protected A-site. As a consequence, the first 4 protected codons of the CDS are mapped to the 5’UTR. In addition, genes with less than 10 cumualtive reads for half of the available samples were removed. The gene list was further filtered on genes containing at least an average 10 RNA-seq reads and an average of 4 ribo-seq reads for all three replicates. This additional filtering step was applied to account for the poly(A) selection, through which transcripts (such as histones) lacking a poly(A) tail are incorrectly identified as significant. Transcripts with a false discovery rate (FDR) < 1% were considered significantly changed. Reported read counts were normalized by counts per million (CPM).

Identification of translation initiation sites (TIS) in Ht treated samples was performed by a previously published bioinformatics peak calling analysis [31]. ORF coordinates were assigned with Mutalyzer [36], In this analysis, peaks were defined as having >40% of all coverage in the first position and a minimum total coverage of 20. Candidate peaks were considered only if they were a maximum distance of 500nt up– or downstream of an annotated coding sequence (CDS). The maximum coverage for the subsequent 5 downstream codons cannot be higher than the candidate peak, and the candidate peak must have at least 10% of coverage relative to the highest candidate to be considered. Statistical analysis of TIS switching was performed using the R package Ime4 (Linear Mixed-Effect Models using ‘Eigen’ and S4) [37], The model was fitted as previously described [31]. Briefly, fixed effects were assigned for location of the TIS location, Tm treatment, and the interaction between the two. Counts were weighted by library size. Significance between models with and without Tm treatment was determined via a chi-squared likelihood-ratio test and corrected via Benjamini-Hochberg (FDR) at a threshold of 5%.

For UCSC browser snapshots we visualised the peak at the first nucleotide of the RFP and the sum of all three replicates. For metagene analysis we used the RiboGalaxy webtool [38],

## Results

### Induction of eIF2 phosphorylation in erythroblasts decreases protein synthesis

To evaluate the effect of eIF2 phosphorylation on mRNA translation in erythroblasts we aimed for a rapid induction of eIF2 phosphorylation that minimalizes secondary effects on mRNA expression, stability or translation. Depletion of iron and heme to activate HRI is a relatively slow process. Average phosphorylation of eIF2 was 2-fold increased upon a 90 minute treatment of erythroblasts with 2.5 μg/ul tunicamycin (Tm) (Fig 1A). Phosphorylation of eIF2 is known to reduce mRNA translation in general [10]. To assess the protein synthesis rate we measured incorporation of the methionine analogue AHA (L-Azidohomoalanine) in erythroblasts during the 90 minute Tm treatment. Alexa Fluor 488, coupled to AHA, was measured in fixed and permeabilised erythroblasts by flow cytometry. Tm treatment reduced de novo protein synthesis by 35% (Fig 1B). To examine whether the reduced protein synthesis rate was due to decreased translation initiation, the polyribosome profile of Tm-treated cells was compared to untreated cells. (Fig 1C). The area under the curve was quantified for 80S and all polysome peaks independently. During Tm treatment the 80S peak and the peak of the first polysome significantly increased (Fig 1D). A shift from heavy towards light polyribosomes and an increase in the 80S monosome peak in Tm-treated cells indicated reduced polysome recruitment. Notably, we do not observe an increase in free ribosomes, rather an accumulation of transcripts with 1 or 2 assembled ribosomes. Together, the results confirm that Tm treatment of erythroblasts induced eIF2 phosphorylation and reduced mRNA translation.

**Fig 1.**
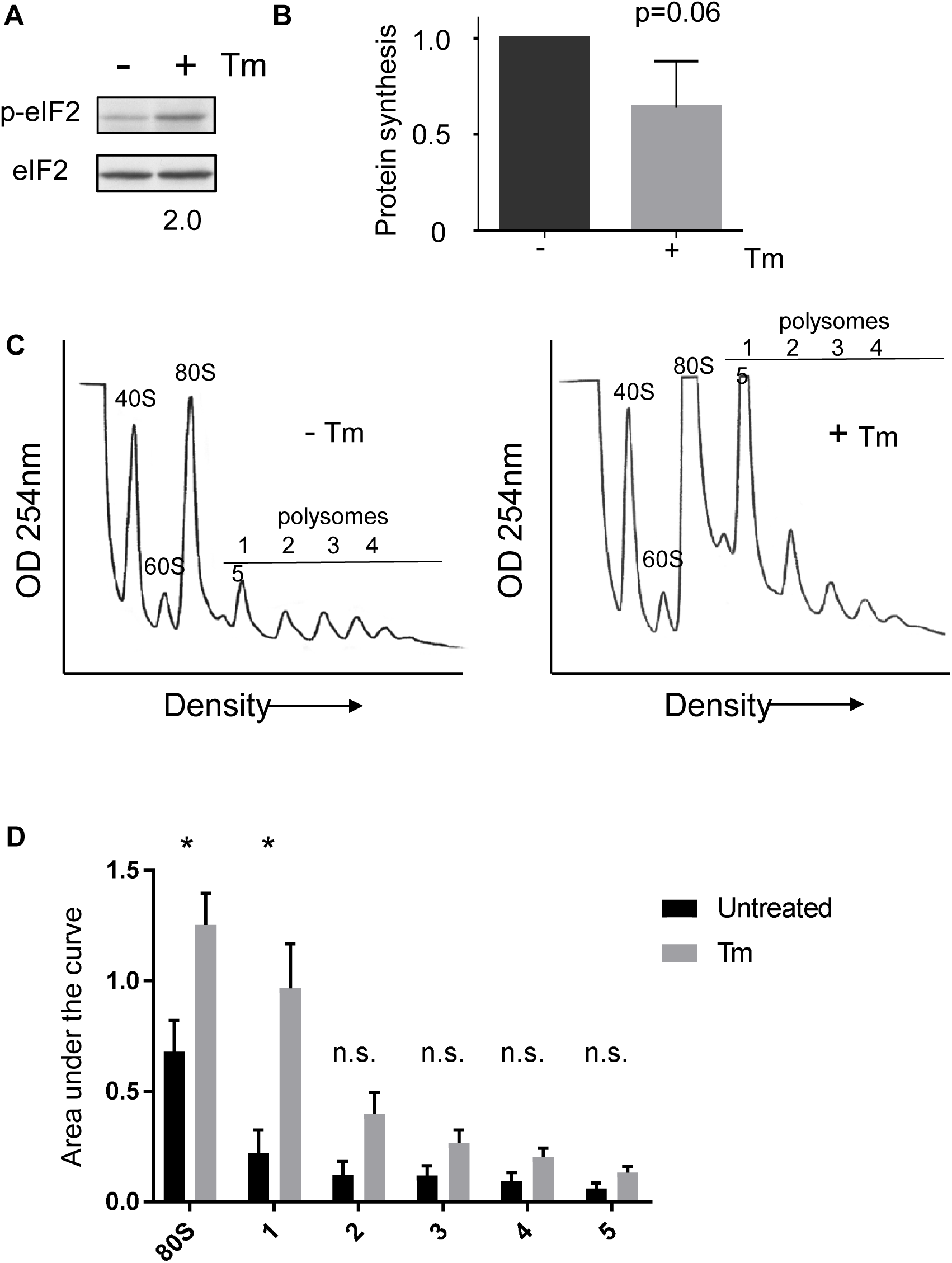
**Phosphorylation of eIF2 reduces protein synthesis.** (A) Murine erythroblasts (line 15.4) were left untreated (-) or treated for 90 min with Tm (2.5 μg/ml). Western blots with total cell lysates were probed for phosphorylated (anti P-S51 eIF2) and total eIF2. Tm increased eIF2 phosphorylation 2-fold (B) Protein synthesis was measured by Click-it technology. Incorporated methionine analogue AHA was coupled to Alexa Fluor 488, and measured by flow cytometry (BD LSR-II). Tm treatment reduces de novo protein synthesis by 35% (average values, n=3, for every pair untreated cells were set to 1, error bar indicated StDev, star indicates p<0.05) (C) Cell lysate was density separated on a 17-50% sucrose gradients and the absorbance at 254nm was measured throughout the gradient, which is a measure for RNA. The polysome profile of untreated cells (left) shows large polysomes with a relatively small monosome peak, whereas Tm-treated cells displayed an accumulation of light polyribosomes (representative plots from 3 independent experiments)

### Tm-induced changes in mRNA translation

To investigate how eIF2 phosphorylation affects translation of individual transcripts in erythroblasts, we compared the ribosome density of transcripts in absence and presence of Tm. For this, ribosome footprint analysis and mRNA sequencing were performed in parallel on 3 biological replicates harvested on separate days. Following 90 min Tm treatment, cells were treated with 100 μg/ml CHX for 5 min to stall elongating ribosomes. Cells were then harvested for ribosome footprint (RFP) and mRNA sequencing analysis. For RFP analysis the cell lysates were treated with RNase-I, after which the resulting monosomes were purified on sucrose gradients, and RNA was isolated. The rRNA fragments were removed with beads, the protected fragments were isolated by PAGE, and library preparation was performed as previously described for myoblasts [31]. The number of reads sequenced per replicate was comparable in all replicates (~15 million, S1 Table). We used STAR to map reads to the genome, because of its capacity to correctly map short reads on either side of an intron. On average, 70-80% of reads mapped to genomic locations, 20-30% of reads were too short and therefore discarded. The modal RFP length was 30-32 nucleotides (S1A Fig). The presence of two populations with distinct footprint length may reflect the two rotating positions of the ribosome and implies that CHX did not completely stall elongation [39], Reads were evenly distributed along all chromosomes, which implied that rRNA fragments were efficiently removed (S1B Fig). CHX stalls ribosomes, but enables preinitiation complexes to assemble and reach the start codon. CHX-induced accumulation of reads at start codons may be enhanced by Tm [40], To investigate whether CHX induced an accumulation of reads at start codons we plotted CHX reads 20 nt upstream or downstream of the start codons of the triplicates separately. This indicated that the majority of the protected fragments start at position −13 (frame 3) from the start codon, instead of the commonly observed position −12 (frame 1). Importantly, CHX reads were similarly distributed along the start codon in Tm-treated and untreated cells (S1C Fig). These results showed that the combined Tm and CHX treatment did not induce severe side effects during stress. Metagene analysis of the protected fragments indicated that the majority of the RFPs are in frame 3 (S1C Fig). Using the same protocol on myoblasts, we previously found frame 1 as the common frame, which may indicate a change in ribosome composition in erythroblasts that makes it difficult to digest the last nucleotide [31,41]. This periodicity is also comparable to previous reports [14].

To use ribosome density as a proxy for protein synthesis in response to Tm-induced eIF2 phosphorylation, we addressed RFPs in the annotated 5’UTR and the protein coding ORF (CDS) separately. RFPs were mapped to the start of the protected fragment at −13 of the P-site. By consequence, the first 4 codons of the CDS mapped to the 5’UTR and are omitted from the analysis of ribosome density on the CDS. We compared the 2Log normalized RFP reads (cpm) of the CDS of all transcripts in Tm-treated erythroblasts to untreated cells (Fig 2A; S2 Table). Ribosome density on the CDS of the classical examples *Atf4* and *Ddit3/Chop* was increased in Tm-treated erythroblasts compared to untreated cells. Transcripts with a more than average reduced ribosome reads due to Tm treatment included, among others, *Mllt1*, *Csde1*, *Dym* and *Pabpc1* (Fig 2A).

**Fig 2.**
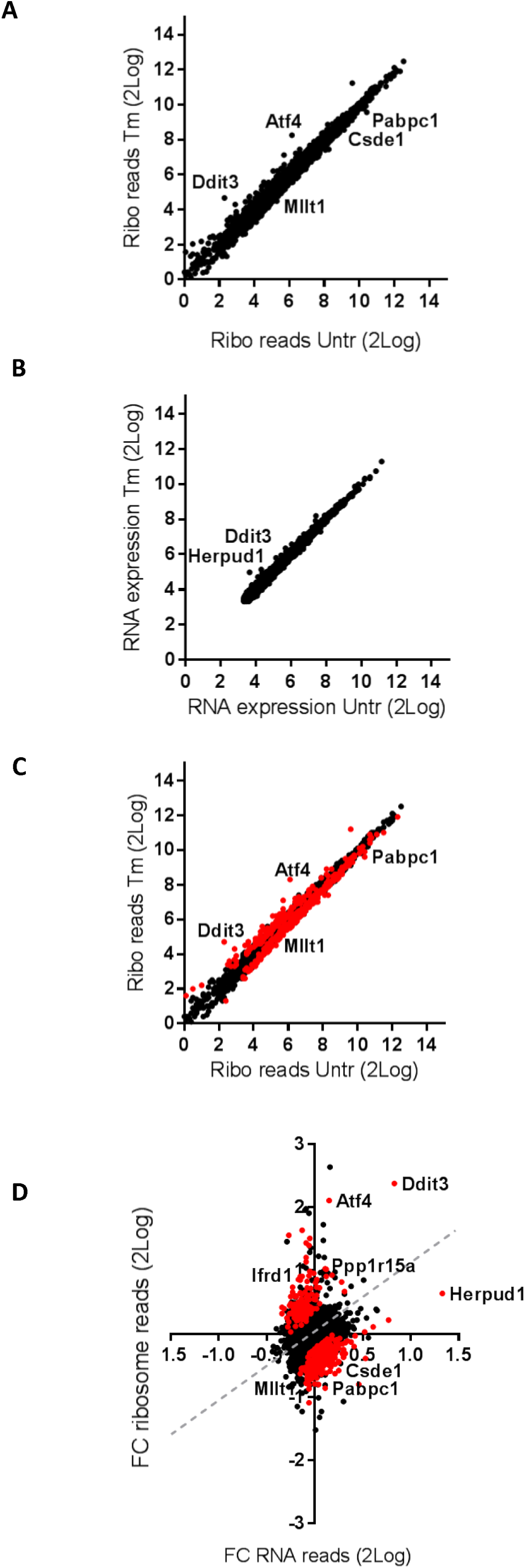
**Tm treatment alters ribosome density on selected transcripts.** (A-B) Murine erythroblasts (line 15.4) were left untreated (-) or treated for 90 min with Tm (2.5 µg/ml). samples were harvested and processed for both ribosome footprinting (A, ribo reads) and polyA+ RNA sequencing (B, RNA expression). Sequence reads of 3 biologically independent experiments were normalized and averaged. For RFP reads we applied a threshold of, on average, 1 read per condition (A), for RNAseq a threshold of, on average, 10 reads per condition (B). For few dots, representing known targets of eIF2 phosphorylation, the transcript name is indicated. (C-D) A statistical interaction model indicated differential ribosome density on 484 transcripts at a FDR <01%, indicated as red dots. Dashed gray line indicates the area where translation follows transcription. (D) The fold-change (FC) in ribo reads (Tm-treated average reads/untreated average reads; Tm/Untr) was plotted against the FC Tm/Untr in RNA expression. Figures are based on data presented in S2 Table.

Tm treatment changes mRNA translation through eIF2 phosphorylation [42], and affects gene transcription through activation of Atf4, Atf6 and Xbp1 [43], To specifically define the effect on mRNA translation, RFP reads must be corrected for mRNA expression. Aliquots of the same cell samples were processed for polyA+ transcriptome analysis. mRNA reads were normalized (cpm), transcripts with an average read intensity <10 cpm were filtered out. The 2Log transformed mRNA reads derived from Tm-treated and control cells were compared. The short Tm treatment hardly induced changes at the RNA level (Fig 2B, S2 Table), although transcription of some genes, among which *Herpud1* and *Ddit3*, was upregulated by Tm.

Combining RFP and mRNA sequencing allows for a more accurate comparison of ribosome density. We employed a statistical model that examines the relationship between RFP and RNA reads (i.e. ribosome density) for each cell sample and calculates the probability that this relation is similar for Tm-treated and control samples (each in triplicate). At a false discovery rate (FDR) of 1%, Tm treatment increased the ribosome reads in 147 transcripts, and decreased the ribosome reads in 337 (Fig 2C; S2 Table). For these transcripts we calculated the fold change (FC) in RFP and in mRNA reads of Tm-treated over control cells from the average cpm (Fig 2C, S2 Table). As expected, Tm treatment increased the translation of *Atf4* and *Ppp1r15a*, with a limited change in transcription. Tm increased *Ddit3* mRNA expression, but also significantly increased its translation (FC increase in RFP significantly higher than in RNA-seq). Other notable translationally upregulated transcripts were *Ibtk* and *Ifrd1/Tis7*. Among the translationally downregulated transcripts during stress were *Csde1* and *Dym*. Interestingly, *Herpud1* stands out because its transcription was increased, whereas its translation rate lagged behind (Fig 2D).

### Pathways that were affected by the Tm treatment

We investigated which pathways were altered by transcripts with significantly altered ribosome density using overrepresentation analysis (ORA) with Genetrail2 [44], Increased ribosome density was foremost associated with transcripts encoding proteins of mitochondria, mitochondrial and endoplasmic reticulum components (enrichment p<10^−6^), followed by transcription complex (p=1.6×l0^-3^) (S3 Table). [45], Among molecular processes, transcriptional (co)activator complexes were most enriched (p=1.3×10^−4^). The stress response factors Atf4 and Ddit3 directly bind DNA to induce transcripts involved in cell survival or apoptosis [43], The transcription factors Gtf2h, Mbd3, JunB and Kmt2e, were also enriched among transcripts with increased ribosome density. For transcripts with more than average decreased ribosome density, the top 30 pathways are shown in S4 Table, according to the adjusted p-value. Among molecular mechanisms, the most enriched transcripts were associated with kinases, and control of kinase activity (p<10^−10^). The second most enriched, and independent molecular function was again transcription activation and chromatin (p=10^−9^). In conclusion, prolonged phosphorylation of eIF2 will reprogram erythroblasts through altered expression of multiple transcription factors, which may stabilise a “stress phenotype” of erythroblasts.

*Detection of translation initiation sites*. In parallel with the CHX treatment, cells were treated with 2μg/ml Harringtonine (Ht) for 7 min to stall initiating ribosomes at start codons, while associated ribosomes complete translation and run off the transcripts. Following quality control, we obtained 11 to 15 million reads per individual sample (triplicate experiments with and without Tm) of which an average of 60% could be mapped to the genome using STAR (S1 Table). We combined STAR with a previously described script that maps the first nucleotide of the RFP and predicts the corresponding translated codon [31]. Similar to the CHX-stabilised RFPs, also the Ht-stalled RFPs mainly started in frame 3 (S2 Fig). Accordingly, most protected reads started at position −13 relative to the annotated start codon (Fig 3A). Because test runs already showed the preferential protection of 13 nt, we had increased the RNAse-l concentration compared to the original protocol that yielded reads starting in >80% at the −12 nucleotide position [31]. This did not make a difference in the length of the pattern of protected fragments. We separated protected fragments according to read length, but longer and smaller fragments were similarly distributed over −12 and −13 (data not shown). Therefore, in our TIS peak detection, we called peaks at both positions.

**Fig 3.**
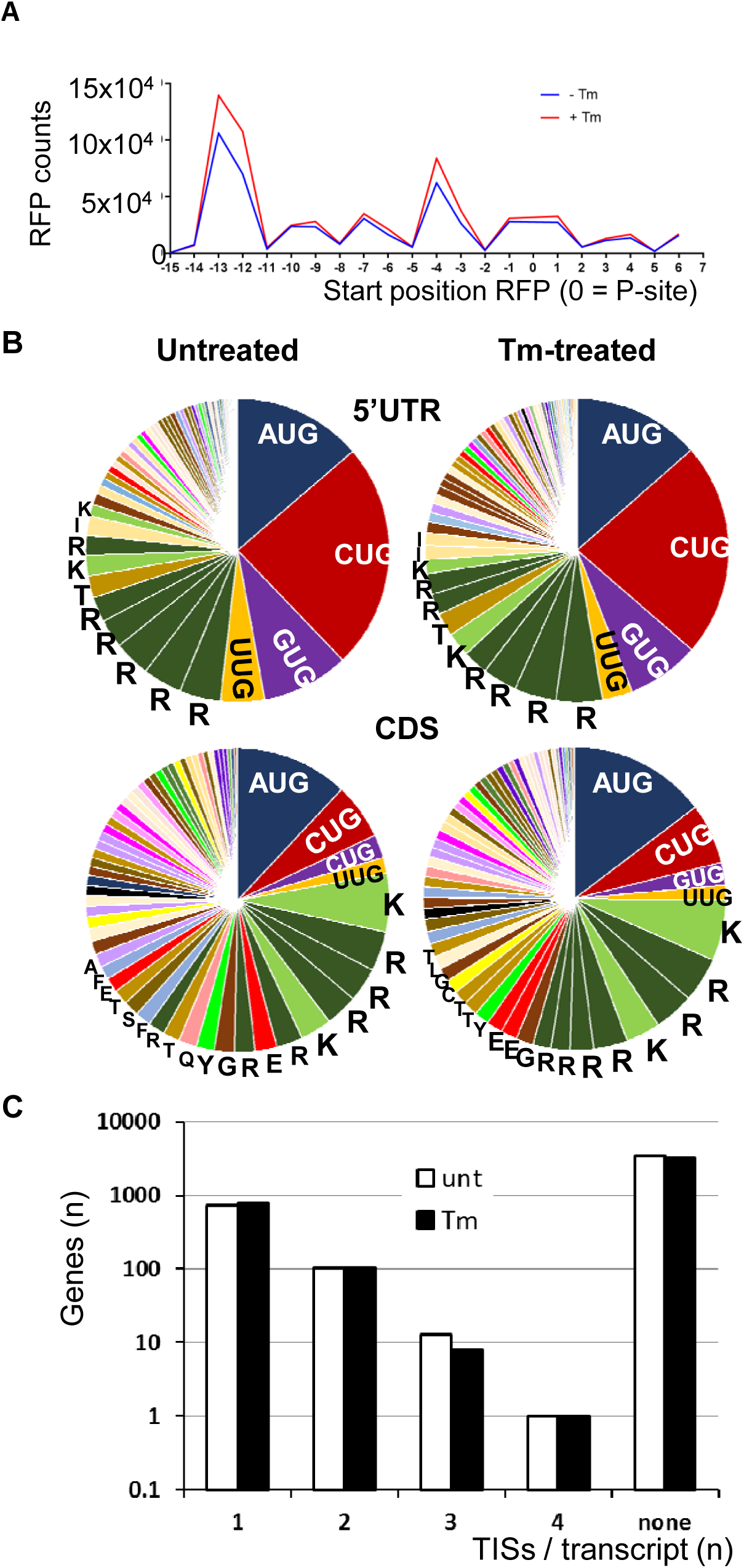
**Characterisation of translation initiation sites (TIS), RFP obtained from cells treated with Harringtonine (Ht; 7 min. 2μg/ml)** (A) The start of the protected RFP fragment, was mapped relative to the the annotated start codon. The start codon is located on position 0, 1, 2 and represents the P-site of the ribosome (because Ht blocks the E-site). The number of RFP reads starting at each position relative to the start codon is indicated (B) mapped Ht RFP were analysed with a peak calling programto define potential TIS in the 5’UTR (top) or CDS (bottom) in cells treated with Tm (right side) or untreated (left side). In the 5’UTR almost half of the detected TIS represented canonical (AUG) and noncanonical (CUG, UUG, GUG) startcodons, whereas only ~25% of all peaks in the CDS represented canonical or noncanonical start codons. The amino acid (1 letter) code of non-start codons was added to the codons that were most frequently detected as putative TIS. Exact percentages and codons are presented in supplemental Table S6. (C) The number of Ht peaks (potential TIS) that were detected in the annotated 5’UTR of individual genes (U: no TIS detected).

The cumulative reads of the triplicate for each condition, as mapped with STAR, were entered into the previously developed peak calling algorithm to identify translation initiation sites (TIS) [31]. Based on their position in the consensus transcript, peaks were segregated to 5’UTR TISs, annotated start codon TISs, TISs in the CDS, or in the 3’UTR. Peak calling was performed both with a setting of peaks at 12nt and at 13nt from the read start. Peaks were assigned to AUG, CUG, GUG or UUG start codons at either +12 or +13 from the start of the protected fragment. All other peaks were assigned to the codon at the +13 position counted from the top of the peak. A total of 1940 and 2175 TISs were identified in the annotated 5’UTRs of transcripts in untreated and Tm treated cells, respectively (S5 Table). From all 5’UTR TISs, 14% were mapped to an AUG codon, 25 and 23% to a CUG codon, 9 and 8% to GUG, and 5 and 3% to UUG in untreated and Tm-treated samples, respectively (Fig 3B, S5 Table). The CDS of untreated and Tm-treated cells revealed 1935 and 2045 TISs, respectively. In the CDS the AUG TISs (12 and 15%) were more abundant than CUG TISs (6 and 7%) (Fig 3B, S6 Table). The preference for CUGs in the 5’UTR, and for AUGs in the CDS is similar to what has been reported [25,31], Overall 53 and 47% of TISs in the 5’UTR were [A/C/G/U]UG startcodons, but these codons only comprised 23 and 26% of all TISs in the CDS. Interestingly, the remainder of the peaks in both the 5’UTR and the CDS was not randomly distributed. Of all TIS peaks in the 5’UTR 24% mapped to triplets encoding the large, and positively charged amino acids Arginine (R) and Lysine (K). In the CDS, 28 and 30% of all peaks mapped to triplets encoding R or K (Fig 3B). To assess whether these are specific artefacts of the Ht treatment, or whether ribosomes pause at these codons, we compared Ht and CHX RFPs in the UCSC web browser. As an example, we show *Abce1* in which we found a Ht peak mapping to an AGG codon, however no CHX reads were present on this location (S3 Fig). The selective presence of the peaks in the Ht track indicated that these are Ht artefacts and not ribosome pausing sites.

### Control of mRNA translation is poorly predicted by uORFs

In the majority of transcripts we detected TISs in the 5′UTR. The 1940 TISs assigned to the 5’UTR of transcripts in untreated erythroblasts corresponded to 1467 genes, as some transcripts carry several mapped TISs. In Tm-treated erythroblasts, 2175 TISs in the annotated 5′UTR corresponded to 1666 genes. However, some of these peaks can be Ht-induced artefacts. Therefore, when we only consider [A/C/G/U]UG start codons as real TISs, TISs were detected in 867 transcripts in untreated erythroblasts and in 907 transcripts in Tm-treated erythroblasts. In most transcripts we detected 1 TIS. The maximum number of detected TISs in the 5′UTR was 4 in the case of *Eri3* (*Exoribonuclease Family Member 3*) (Fig 3C, S5 Table). Taken together this means that uORF translation is widespread among expressed genes in both conditions.

In theory, comparison of TIS peak intensities corresponding to annotated start codons should validate the differences in ribosome density. Increased or reduced ribosome density should be mirrored by increased or reduced peak hight on the start codon. However, start sites hardly accumulate reads when they are located downstream of an uORF, and the division of the peak over the −12 and −13 position also complicated quantitative analysis. The analysis of ribosome density was much more accurate than an analysis of peaks on annotated start sites. Therefore we focussed in the presence of unexpected start sites within the CDS that may give rise to proteins with distinct N-termini. We considered all genes with at least 1 observed [A/C/G/U]UG consensus start codon TIS in the 5’ UTR. For 683 genes we identified consensus start codon TISs under both control and Tm-treated conditions (Fig 4A, blue, bold numbers). The high overlap (79% of the lowest number) is expected, because the first TIS peak accumulates during Ht treatment while the formation of pre-initiation scanning complexes and scanning from the cap continues. For TISs in the CDS we also only considered [A/C/G/U]UG TISs. TISs in the protein coding domain are often an underestimation, because stalling of ribosomes at subsequent start codons depends on scanning complexes that had passed the initial TIS at the start of Ht treatment. Among the 683 transcripts with a TIS detected in the 5′UTR of transcripts from both Tm-treated and untreated erythroblasts, we detected a TIS in the CDS of 41 transcripts: 21 TISs in the CDS of transcripts of both TM-treated and untreated condition, 12 TISs only in the transcripts of Tm-treated cells, and 8 TISs only in transcripts of control cells. (Fig 4A).

**Fig 4.**
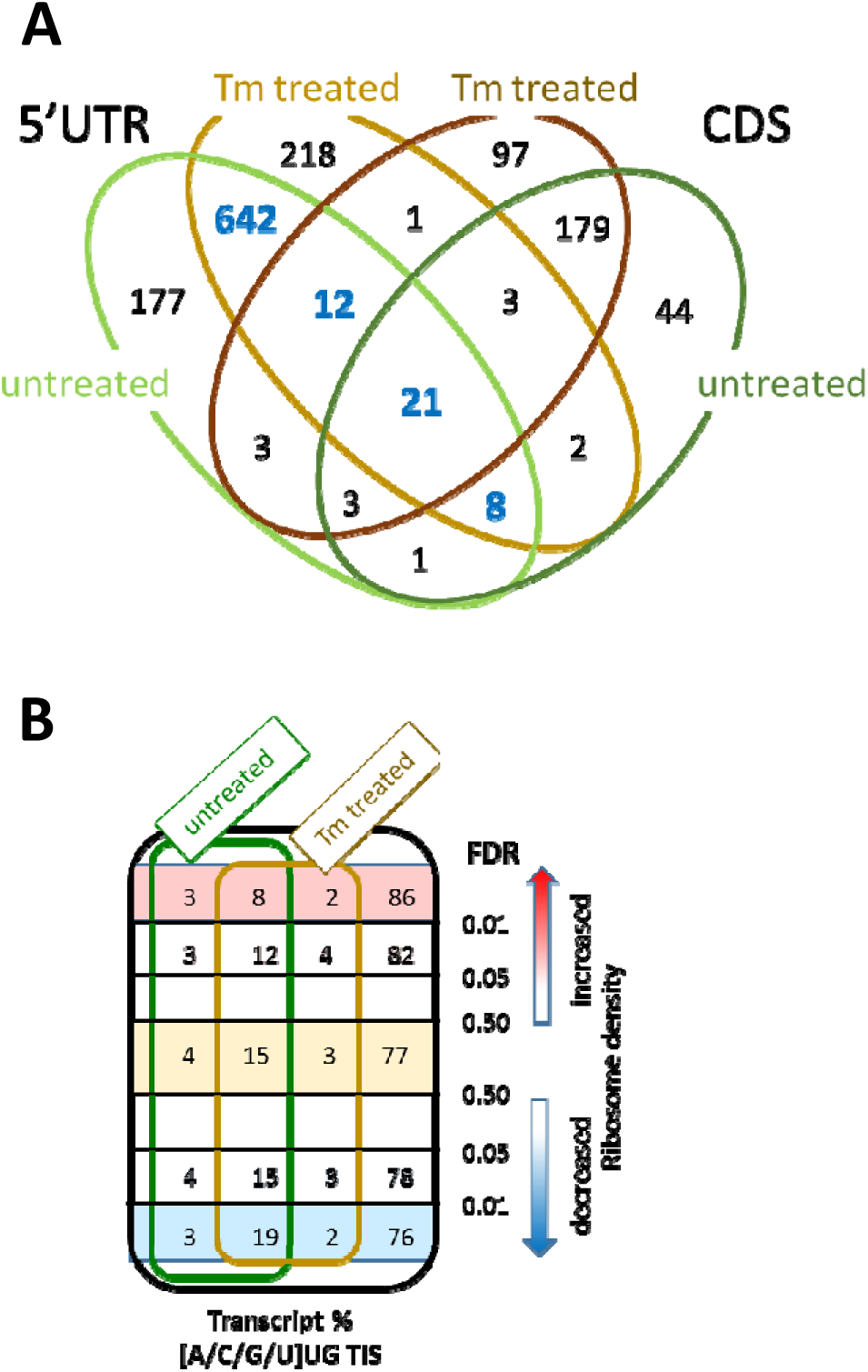
**Distribution of TIS in the 5’UTR and CDS of transcripts dependent on Tm treatment.** (A) Venndiagram showing transcripts with TIS detected in untreated cells (green), and/or Tm-treated cells (brown), and detected in the 5’UTR (light color, any TIS) or in the CDS (darker color; start codons only (AUG/CUG/UUG/GUG)). Bold blue numbers are transcripts with at least one predicted TIS in the 5’UTR under both conditions; predicted TIS in the CDS of these transcripts may give rise to alternative proteins dependent on Tm treatment. (B) total transcripts (black circle) and transcripts with TIS detected in the 5’UTR of untreated (green circle), and/or Tm-treated cells (brown circle) with Tm-induced increased ribosome density (red squares on top, FDR<0.01 or <0.05); similar ribosome density (orange, FDR>0.5); or reduced ribosome density (blue squares on bottom; FDR<0.01 or FDR<0.05). Both transcript numbers and percentage are indicated.

A detected peak in the coding sequence may indicate translation of an ORF that leads to a protein isoform. An example is *Transcription factor cp2* (*Tfcp2*) which is translated from the annotated start codon embedded in a strong Kozak consensus sequence. A second very strong TIS peak maps downstream of the start codon in the CDS. However, it does not correspond to a N-terminally truncated protein but to a 9 codon small ORF (S4 Fig), which appeared to be the case for more peaks in the CDS.

To assess which TISs are actually affected by Tm exposure, we investigated whether Tm treatment changed the peak intensity ratio between TIS peaks within a transcript as previously described [31]. The ratio between triplicate TIS peak reads at distinct positions within a gene was compared between untreated and Tm-treated cells. At a p-value less than 0.01 few transcripts revealed differentially employed TISs in their 5’UTR (S7 Table). For example, the ratio between the TIS detected in the 5’UTR of *Ranbp1* and the TIS of the annotated CDS start codon differed significantly dependent on Tm treatment (S5 Fig). Interestingly, *Ranbp1* RNA expression in erythroid progenitors is high compared to CD34+ cells [46], In conclusion, we did not detect major changes in the expression of protein isoforms upon phosphorylation of eIF2.

Next we investigated the role of uORFs in the quantitative control of RNA translation. We hypothesized that uORFs render translation of the protein coding ORF more sensitive to eIF2 phosphorylation. To assess whether increased, or decreased ribosome density in the CDS upon eIF2 phosphorylation is due to uORF translation, we considered transcripts with at least 1 detected TIS peak.

We divided transcripts with at least 1 TIS, into pools based on i) FDR interaction term significance for translation efficiency in response to Tm treatment and ii) whether ribosome density was increased or reduced (S2 Table, Fig 4B). As a control group of transcripts that were not specifically affected by eIF2 phosphorylation, we considered transcripts with a FDR>0.5 and at least 1 [A/C/G/U]UG TIS detected in absence and presence of Tm. In this group 77% of transcripts was without TIS, and 15% harboured at least 1 TIS in the 5’UTR both in absence and presence of Tm (Fig 4B). Surprisingly, this distribution of transcripts with or without TISs in the 5’UTR was the same for the genes in which ribosome density was increased (Pearson′s Chi-square p-value not significant). Among the transcripts with increased ribosome density at a FDR<0.05 19% contained a TIS (12% under both conditions), and among transcripts with more than average decreased ribosome density (FDR<0.05) 22% contained a TIS (15% under both conditions) (Fig 4B). These results suggest that translation of an uORF does not seem to be a strong predictor of either quantitative or qualitative mRNA translation.

### Long uORFs with a CUG start codon occur commonly in transcripts with Tm-enhanced translation

For individual transcripts, the translation of uORFs can be crucial for proper regulation. For transcripts of which translation was up- or more than average downregulated in response to Tm treatment, we established the TIS positions (Ht-induced TIS peaks) and the sizes of corresponding uORFs (RFPs protected in presence of CHX) (Fig 5). We first analysed 10 transcripts with Tm-increased ribosome density and upstream TISs (Fig 5A). We detected 14 TIS in the 5’UTR of these 10 transcripts: 2 UGU, 6 CUG and 6 AUG codons. From the 6 AUG codons 4 mapped to the known targets *Atf4*, *Ddit3* and *Ifrd1*. Thus, the novel, experimentally determined TISs were mainly non AUG. These non-AUG TISs that we established experimentally are hard to predict, particularly when they occur in a poor Kozak consensus sequence (e.g. the Cag CUG C start codon in *Mbd3*).

**Fig 5.**
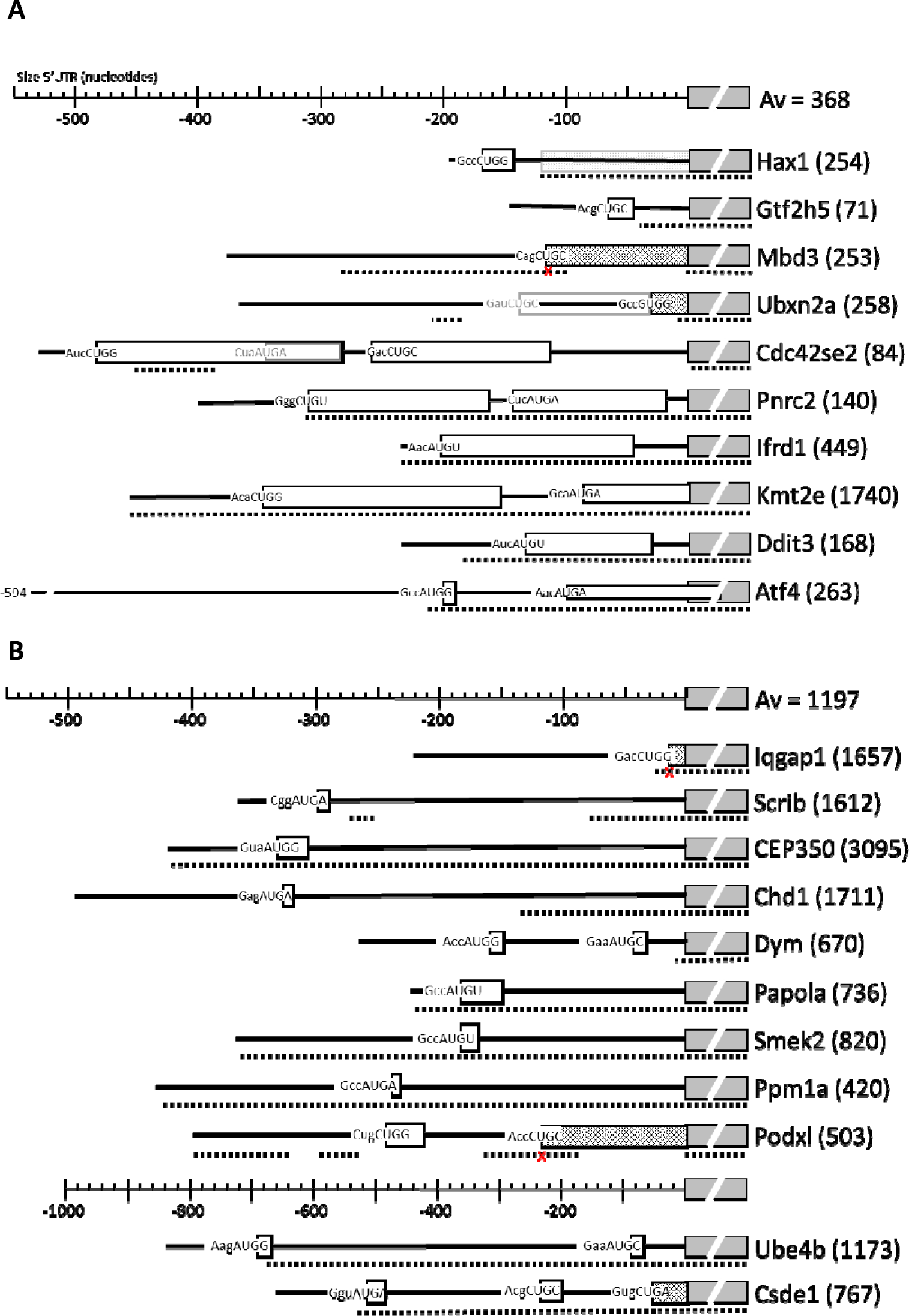
**Position and length of uORFs in the 5’UTR of transcripts subject to Tm-controlled translation.** (A, B) Top line indicates the distance in nt upstream of the main annotated start codon. The same relative size is used for transcripts with Tm-enhanced translation (A), or Tm-decreased translation (B), except for the last two transcripts for which size was condensed 2-fold as indicated by a separate size marker. The collapsed annotated protein ORF is shown by a grey interupted box, with the protein name directly at the rightside. Numbers between brackets indicate the size of the main annotated protein in amino acids. Adjacent boxes with a fence pattern on the left of the “protein box" indicate a N-terminal extension of the protein. A fenced box at the back ground as shown for Hax1 indicates that this part of the annotated protein seems not translated. All uORF are indicated by open boxes, and the start codon is written at the start of the box including its Kozak context. The dashed line below indicates areas that are >90% conserved between mouse and man. A small cross below the start codon in a conserved area indicates that the start codon is not conserved. Conserved areas were identified by Blastn with the mouse sequence on the human transcriptome.

The mechanism employed by *Atf4*, a small uORF followed by an inhibitory uORF overlapping the protein codon TIS, appeared unique for *Atf4*. In only two other transcripts small uORFs were translated *(Hax1* and *Gtf2h5*), and in only one transcript a second uORF overlapping the protein start codon was translated *(Kmt2e*). Strikingly, the annotated start codon of *Hax1* was skipped, and an AUG codon 120 nt downstream was used as the TIS for *Hax1* coding frame. The GWIPS website (http://gwips.ucc.ie/) [47] revealed that this occurs in most mouse cells. The novel TISs in *Mbd3* and *Ubxn2a* appeared to be in frame with the known CDS and initiated an N-terminally extended protein isoform. Comparison with global data on the GWIPS website indicated that this is common for *Mbd3* in mouse cells. In contrast, most cell types are protected from the extension of *Ubxn2a* by a large uORF that ends just 1 codon upstream of the TIS. This uORF was hardly expressed in erythroblasts according to both Ht- and CHX-induced RFPs. The N-terminally extended isoforms of *Mbd3* and *Ubxn2a* are not conserved between mouse and human.

In five transcripts one or two long uORFs were translated, four of these are >90% conserved between mouse and human. These uORFs are also translated in other celltypes (GWIPS data), although at different ratios. Strikingly, an AUG codon within the first long uORF of *Cdc42se2* is the major TIS detected in most other cells. In our data this was a minor start, and we found a major contribution of the two long uORFs, both in Ht- and in CHX-arrested RFPs. The TIS of the first uORF of *Pnrc2* is hardly detected in other cell types (GWIPS). In absence of Tm, we detected less CHX-stabilised RFP in the first uORF compared to the second uORF (S6 Fig). Tm, that induced a 1.9-fold increase in *Pnrc2* translation, changed the relative density of ribosomes on the two uORFs in favour of uORF1, which is even more distant from the global aggregate on the GWIPS site.

### A long 5’UTR with a short uORF harbouring an AUG TIS is common in transcripts undergoing Tm-reduced translation

The start codon, length, and position of uORFs in transcripts with more than average Tm-decreased translation was different from the uORFs found in upregulated transcripts (Fig 5). Whereas we detected many long uORFs in transcripts with Tm-enhanced translation, all uORFs detected in transcripts with Tm-reduced translation are short. In 11 transcripts (>2-fold reduction compared to average) we observed 15 TISs, 11 of which were AUG codons. For 3 of the 11 transcripts we observed an N-terminal extension (*Iqgap1*, *Podxl*, *Csde1*), that are also observed for other cell types but in a lower frequency (GWIPS comparison). The uORF of *Chd1* is not detected in other cell types, whereas an additional, further upstream, uORF was detected for *Ppm1a* and *Csde1* in many other cell types, but not in our erythroblasts (GWIPS comparison). The 5’UTR of seven transcripts is >90% conserved between mouse and man, suggesting conserved mechanism of translation control. Notably, 9/10 transcripts subject to Tm-enhanced translation encoded short proteins (average of all encoded proteins is 368 amino acids). In contrast, the average of protein size encoded by transcripts subject to Tm-decreased translation is 1197 amino acids.

## Discussion

Iron deficiency, oxidative stress, or the presence of unfolded proteins in erythroblasts activates the eIF2 kinases HRI and PERK, respectively, which results in phosphorylation, and thereby inactivation, of eIF2. This decreases overall mRNA translation to prevent for instance the accumulation and aggregation of globin polypeptides in absence of iron and heme [9], To characterise the molecular pathways and cellular processes that respond to eIF2 phosphorylation in erythroblasts we combined ribosome profiling and transcriptome analysis to detect transcripts with increased ribosome density, or with a more than average decreased density of elongating ribosomes upon eIF2 phosphorylation. We found, among others, known components of the ISR pathway to be increased in translation, such as *Atf4, Ddit3*, and *Ifrd1*, but also transcripts that are less well known to be translated upon eIF2 phosphorylation including *Prnc2*, that encodes a protein involved in recruitment of transcripts to P-bodies for subsequent degradation [48], On the other hand, stress also led to more than average downregulation of translation for a set of transcripts that included *Csde1* and *Dym*. Whereas stabilisation with CHX identified elongating footprints, the treatment of erythroblasts with Ht identified footprints at translation initiation sites. Combination of CHX and Ht RFPs showed that the presence of a translated uORF did not predict the sensitivity of a particular mRNA during eIF2 phosphorylation. The high degree of conservation between the 5′UTR of man and mouse suggests that the translation mechanism may be more complex than only the presence of uORFs. Strikingly, transcripts with Tm-enhanced translation contained long, conserved uORFs that often started with a CUG start codon, whereas transcripts with Tm-reduced translation contained short uORFs starting from an AUG codon.

Some of the transcripts that we found to be translationally upregulated upon Tm treatment of erythroblasts were recently linked to eIF2 phosphorylation in HEK293 cells. These transcripts encoded proteins involved in the ISR such as Atf4, Atf5, and Ppp1R15a/Gadd34, Ibtk, and Tis7/IFRD1 [14,49], The ISR is highly conserved between eukaryotes, from yeast to mammals [50], Several ribosome profiling datasets were published that address the ISR, but the data are difficult to compare. Lack of uniformity in methods, in induction of eIF2 phosphorylation, in statistical analysis and in cell types complicates comparisons between these studies. Nevertheless, we compared the transcripts with increased translation in erythroblasts to transcripts with increased ribosome density in response to arsenite treatment of HEK293 cells [14]. Whereas we (*i*) identified differential ribosome density in erythroblasts, and (*ii*) used a statistical interaction model to compare RFP and RNAseq reads. Andreev et al. (*i*) calculated translation efficiency in HEK293 cells, and (*ii*) determined the Z-score for the fold-change in translation efficiency. They considered transcripts with a Z-score>4 as significantly upregulated. For this comparison we considered the transcripts with a Z-scoreώ3 in the dataset of Andreev et al. (S7 Fig). Strikingly, the overlap between differentially translated transcripts was limited to *Atf4, Atf5, PpplRlSa, Slc35A4* and *Ifrd1*. There was a clear separation between transcripts that were differentially translated in HEK293 cells or in erythroblasts. This difference seems too large to be due to technical differences and may rather reflect an essential difference between these two cell types. This implies that the ISR downstream of eIF2 phosphorylation is different in erythroblasts compared to HEK293 cells. The activity and specificity of eIF2 is modulated by the association with eIF1 and eIF5 [51]. eIF1 is upregulated in response to SCF-induced erythroblast expansion, whereas eIF5 is upregulated during differentiation to hemoglobinised, enucleated red blood cells [22], Interestingly, cancer cells were also shown to modify their response to eIF2 phosphorylation by expression of the alternative translation initiation factor eIF2A [52], The effect of eIF2A only becomes apparent when eIF2 is limiting [53], Thus, depending on the expression levels of various translation initiation factors, each cell may respond differently to eIF2 posphorylation, because translation of uORFs and protein coding ORFs will depend on the combination of eIF2 availability plus the modulation of its activity and specificity by associated initiation factors.

Interpretation of RFP data sets, and particularly of translation initiation sites is complicated by several factors including (*i*) sequence depth, (*ii*) ligation bias, and (*iii*) TIS peak imbalance. First, each read is a single count on a single codon. A substantial number of reads need to map to each codon position to identify changes in codon usage that are statistically significant. From samples treated with CHX we obtained a total of >45 million reads for the combined triplicate. Statistical analysis uses the individual experiments. Thus, peaks that can be discerned in the UCSC web browser may still lack statistical power. Second, we observed that ligation of the small RFP fragments to adapter oligonucleotides is very sensitive to bias and that this bias depends on the ligation kit. We detected the start codon of the first uORF of *Atf4* in pilot experiments, but the final experiment only showed a relatively low number of reads at this position. We cannot exclude the possibility that the use of a different adapter ligation kit introduced bias in the ligation step. In agreement with this supposition, ribo-seq profiles of *Atf4* also show a loss of uORF1 in other studies that used the same library prep kit [54,55] compared to studies that use different methods, as shown in the GWIPS-viz genome browser [47], Third, the detection of TISs following Ht treatment has a strong bias towards the most upstream uORF. Ht or CHX do not inhibit the association of the pre-initiation scanning complex at the cap, and scanning to the first start codon. During treatment, this first peak continues to grow, while all other peaks downstream of the first peak depend on scanning complexes present between the peaks at the start of the treatment.

Finally, we also observed an enrichment of Ht peaks at codons that code for Arginine (R) and Lysine (K). These amino acids are positively charged, and they are among the bulkiest amino acids. The triplets coding for other bulky amino acids (tyrosine, Y; Phenylalanine, F) are not enriched among the peaks. Having a positively charged (large) amino acid at the P-site of the ribosome may either create more space at the A-site to bind Ht, or it may pause ribosome progression. In the latter case ribosome density should also be increased upon CHX treatment. Therefore, TIS peaks are subject to bias and need to be interpreted with caution. In combination with elongating RFPs, however, it is a powerful method to identify uORFs. Ribosome profiling on other cell types reported different biases [25,56], This may be due to technical details such as bias in the isolation and ligation of protected fragments, but it could also hint at a cell type specific composition of the pre-initiation scanning complex and elongating ribosomes.

The data also show that many alternative start codons, particularly CUG, are used as TISs. Therefore, prediction of uORF translation from the primary transcript sequence is difficult, if not impossible. Experimental TIS analysis such as the Ht treatment to stall ribosomes at start codons, is needed to understand how TIS may contribute to control translation in specific transcripts. Selective translational control by eIF2 is performed through differential start codon recognition and the presence of uORFs on 5’ UTRs of specific mRNAs [8], However, in our proteotoxic stress model we did not find an enrichment of uORF containing transcripts. The translation of uORFs appeared widespread.

The transcripts with significantly altered translation compared to the average change in translation caused by Tm were enriched for CDS giving rise to transcription factors, like Pnrc2, Tis7, Kmt2e and JunB. Pnrc2 interacts with the glucocorticoid receptor to induce mRNA decay of some transcripts [57], Glucocorticoids are important for expansion of the erythroblast compartment upon induction of stress erythropoieses [58], Interestingly, JunB was reported to drive erythroid differentiation [59], Increased expression of JunB in response to eIF2 phosphorylation may be a convergence node in erythropoiesis for ER-stress and activation of stress kinases of the MAPkinase pathway similar to what was found for pancreatic cells [60], Tis7 was found to be upregulated in chicken erythroid cells during hypoxic stress [61]. Kmt2e regulates cell cycle progression in myoblasts [62], These transcription factors could also be involved in activating the transcription of other stress responsive genes and induce a cell survival mechanism in erythroblasts.

In conclusion, translational control by eIF2 in erythroid cells is important for maintaining red blood cell function and survival. In this study we have used ribosome profiling to investigate which transcripts are translationally up or downregulated during ER stress in erythroblasts. Unexpectedly, uORFs are not enriched in these transcripts. We also observed [A/C/G/U]UG TISs within the CDS of 179 transcripts, and these were mostly short out-of-frame ORFs. Whether these are unimportant side effects due to leaky scanning of the CDS starting codon, whether their translation interferes with the translation of the CDS, or whether the encoded peptides are stable is not known and needs to be investigated. Future studies should be performed to gain more insight into control of translation by eIF2, and to understand the role of these encoded proteins in erythropoiesis.

## Accession numbers

Original sequencing results have been deposited in the BioProject Database under project ID PRJNA380970.

### Data access for reviewers

UCSC browser session:

https://genome.ucsc.edu/cgi-bin/hgTracks?hgS_doOtherUser=submit&hgS_otherUserName=ksmll3&hgS_otherUserSessionName=TIS%20lfrdl%20Har%20%26%20Chx

SubmissionID: SUB2489513

BioProject ID: PRJNA380970

BioSample accessions: SAMN06660139, SAMN06660140

http://www.ncbi.nlm.nih.gov/biosample/6660139

http://www.ncbi.nlm.nih.gov/biosample/6660140

## Acknowledgements

We want to thank Dr E. van den Akker for critical reading of the manuscript, Drs Henk Buermans and Yavuz Ariyurek, Leiden Genome Technology Centre (LGTC), Leids Universitair Medical Centre (LUMC), for deep sequencing support.

## Supporting information

**S1 Fig. Ribosome profiling data quality.** (A) Ribosomes were stabilised with CHX. Shown is the fitted line through the average values of three biological replicates harvested following Tm treatment or three control replicates. Error bars indicate standard deviation. (B) RFP fragments were mapped to the genome and the number of reads (all experiments combined) was enumerated per chromosome. Shown is the percentage of all reads mapping to the different chromosomes. (C) RFP sequence data were uploaded to the RiboGalaxy webtool. The start of each RFP was mapped to the genome. The number of reads starting at position −20 to +50 compared to the startcodon, and on position −50 to +20 compared to the stopcodon were calculated for reads of 32 nt. Reads in each frame are indicated by distinct colors. Red: frame 1, green: frame 2, blue: frame 3. Representative plots of one replicate of each condition is shown.

**S2 Fig. Harringtonine-induced RFP are mostly translated in frame 3.** We used STAR to map Ht-stabilized RFP to the genome, and used our previously described script to map the first nucleotide relative to the annotated reading frame. Shades of blue (a2, b2, c2) represent RFP from untreated cells, shades of orange (a4, b4, c4) represent RFP from Tm-treated cells.

**S3 Fig. Web browser snapshot of *ATP-binding cassette sub-family E member 1 (Abce1)*.** Cumulative Ht- and CHX-stabilized RFP counts from Tm-treated and untreated (Unt) cells are mapped to the genome and visualized in the UCSC web browser. Numbers on the right hand side indicate maximum read counts in the respective lane. Gray lines indicate introns. The arrow indicates a peak of Ht-stabilised RFP that corresponds to a non-start codon. This peak is not present in CHX-stabilised RFP, indicating that this is most likely a Ht-induced artefact.

**S4 Fig. Web browser snapshot of *Tfcp2*.** Aggregate Ht- and CHX-stabilized RFP reads from Tm-treated and untreated (Unt) cells are mapped to the genome and visualized in the UCSC web browser. Numbers on the right hand side indicate maximum read counts in the respective lane. Arrows indicate Ht peaks. Gray lines indicate introns. Part of the 3’UTR is cropped.

**S5 Fig. Web browser snapshot of *Ranbp1*.** Cumulative Ht- and CHX-stabilized RFP counts from Tm-treated and untreated (Unt) cells are mapped to the genome and visualized in the UCSC web browser. Numbers on the right hand side indicate maximum read counts in the respective lane. Gray lines indicate introns. The uORFs in the 5’UTR and the protein coding ORF (CDS) are indicated.

**S6 Fig. Web browser snapshot of the 5’UTR of *Proline Rich Receptor Coactivator 2 (Pnrc2)*.** Aggregate Ht- and CHX-stabilized RFP counts from three replicates of Tm-treated and untreated (Unt) cells are mapped to the genome and visualized in the UCSC web browser. Numbers on the right hand side indicate maximum read counts in the respective lane. Introns have been skipped. The data indicate two uORF that are depicted by grey boxes. Only the start of the protein coding ORF is shown. Two arrows in the top lane (Unt, Ht RFP) indicate TIS at start codons (CUG for uORF1; AUG for uORF2). uORF1 is 48 codons in length, uORF2 is 42 codons in length, they are separated by 17 nt, and the distance between uORF2 and the AUG start codon is 14 nt.

**S7 Fig. Comparison of ribosome occupancy in response to eIF2 phosphorylation in HEK293 cells (Andreev et al.) and mouse erythroblasts (Paolini et al.).** Triangles indicate transcripts of which translation is similarly upregulated upon eIF2 phosphorylation in both studies. White circles represent transcripts with enhanced translation (Z-score >3) in HEK293 cells but not in mouse erythroblasts; dark grey circles represent transcripts with enhanced translation in mouse erythroblasts (FDR<0.01) but not in HEK293.

**S1 Table. Overview of ribosome footprint reads mapped with STAR.** Ribosome reads were mapped with STAR to the genome. This table gives an overview of read length and how many reads mapped to the genome for each sample. Note: Multi-mapped reads were not discarded, unless they mapped to more than 20 locations.

**S2 Table. Normalised sequence counts for ribosome footprints (RFP) and pA+ RNA sequencing (counts per million; cpm).** 2Log normalized RFP reads (cpm) of the CDS of all transcripts in Tm-treated cells were compared to untreated cells. List of significantly altered transcripts during Tm treatment in erythroblasts, cpm values are given foreach sample for ribosome profiling and RNAseq.

**S3 Table. List of upregulated transcripts during Tm treatment.** Upregulated targets were uploaded on Genetrail2 to investigate enrichment of cellular component, biological processes and molecular function.

**S4 Table. List of downregulated transcripts during Tm treatment.** Downregulated targets were uploaded on Genetrail2 to investigate enrichment of cellular component, biological processes and molecular function.

**S5 Table. Translation initiation sites detected by stalling of ribosomes in the presence of Harringtonine.** Peaks were called with the cumulative reads of each triplicate, with our previously developed peak calling algorithm to identify translation initiation sites (TIS). Peaks were divided into 5’UTR TISs, annotated start codon TISs, TISs in the CDS, or in the 3’UTR. The analysis was performed both with a setting of peaks at −12nt and at −13nt from the read start. Peaks were assigned to AUG, CUG, GUG or UUG start codons at either +12 or +13 from the start of the protected fragment. All other peaks were assigned to the codon at the +13 position counted from the top of the peak. TISs in the 5’UTR, the CDS, annotated starts were fused to gene name in random order. Positions are +13 positions, unless a atg, ctg, gtg or ttg occurs at +12, or +14. in that case the atg, ctg, gtg or ttg was preferred.

**S6 Table. Codons at −13 (P) position of translation initiation sites, measured after ribosome stalling with Harringtonine.** Called peaks and triplet codons were compared in untreated and Tm-treated erythroblasts.

**S7 Table. Transcripts with differential use of TIS in absence and presence of tunicamycin.** Peak intensity ratio between TIS peaks in stressed cells were compared to untreated cells for specific transcripts. At a p-value less than 0.01 few transcripts revealed differentially employed TISs in their 5’UTR Coverage: cumulative reads of the peak. Statistics: two way ANOVA between triplicate samples of both conditions

## References

1. Gozzelino R, Arosio P. Iron Homeostasis in Health and Disease. Int J Mol Sci. 2016;17: 1–14. doi:10.3390/ijms17010130

2. Chung J, Chen C, Paw BH. Heme metabolism and erythropoiesis. Curr Opin Hematol. 2012;19: 156–162. doi:10.1097/M0H.0b013e328351c48b

3. Chen J-J. Regulation of protein synthesis by the heme-regulated eIF2alpha kinase: relevance to anemias. Blood. 2007;109: 2693–9. doi:10.1182/blood-2006-08-041830

4. Kühn LC. Iron regulatory proteins and their role in controlling iron metabolism. Metallomics. 2015;7: 232–243. doi:10.1039/C4MT00164H

5. Horvathova M, Kapralova K, Zidova Z, Dolezal D, Pospisilova D, Divoky V. Erythropoietin-driven signaling ameliorates the survival defect of DMT 1-mutant erythroid progenitors and erythroblasts.Haematologica. 2012;97: 1480–1488. doi:10.3324/haematol.2011.059550

6. Bandyopadhyay S, Brittenham GM, Francis RO, Zimring JC, Hod EA, Spitalnik SL. Iron-deficient erythropoiesis in blood donors and red blood cell recovery after transfusion: initial studies with a mouse model. Blood Transfus. 2017;15: 158–164. doi:10.2450/2017.0349-16

7. Lu L, Han A, Chen J. Translation Initiation Control by Heme-Regulated Eukaryotic Initiation Factor 2 α Kinase in Erythroid Cells under Cytoplasmic Stresses Translation Initiation Control by Heme-Regulated Eukaryotic Initiation Factor 2 α Kinase in Erythroid Cells under Cytopl. Mol Cell Biol. 2001;21: 7971–7980. doi:10.1128/MCB.21.23.7971

8. Hinnebusch AG. Molecular mechanism of scanning and start codon selection in eukaryotes. Microbiol Mol Biol Rev. 2011;75: 434–67. doi:10.1128/MMBR.00008–11

9. Chen J-J. Translational control by heme-regulted eIF2α kinase during erythropoiesis. Curr Opin Hematol. 2014;21: 172–8. doi:10.1097/M0H.0000000000000030

10. Wek RC, Jiang H-Y, Anthony TG. Coping with stress: eIF2 kinases and translational control. Biochem Soc. 2006;34: 7–11. doi:10.1042/BST20060007

11. Vattem KM, Wek RC. Reinitiation involving upstream ORFs regulates ATF4 mRNA translation in mammalian cells. PNAS. 2004;101: 11269–11274.

12. Lu PD, Harding HP, Ron D. Translation reinitiation at alternative open reading frames regulates gene expression in an integrated stress response. J Cell Biol. 2004;167: 27–33. doi: 10.1083/jcb.200408003

13. Palam LR, Baird TD, Wek RC. Phosphorylation of eIF2 facilitates ribosomal bypass of an inhibitory upstream ORF to enhance CHOP translation. J Biol Chem. 2011;286: 10939–49. doi:10.1074/jbc.M110.216093

14. Andreev DE, O’Connor PB, Fahey C, Kenny EM, Terenin IM, Dmitriev SE, et al. Translation of 5’ leaders is pervasive in genes resistant to eIF2 repression. Elife. 2015;4: 1–21. doi:10.7554/eLife.03971

15. Calkhoven CF, Muller C, Martin R, Krosl G, Pietsch H, Hoang T, et al. Translational control of SCL-isoform expression in hematopoietic lineage choice. Genes Dev. 2003;17: 959–64. doi:10.1101/gad.251903

16. Suragani RNVS, Zachariah RS, Velazquez JG, Liu S, Sun C-W, Townes TM, et al. Heme-regulated eIF2α kinase activated Atf4 signaling pathway in oxidative stress and erythropoiesis. Blood. 2012; 119: 5276–84. doi:10.1182/blood-2011-10-388132

17. Masuoka HC, Townes TM. Targeted disruption of the activating transcription factor 4 gene results in severe fetal anemia in mice. Blood. 2002;99: 736–745. doi:10.1182/blood.V99.3.736

18. Jousse C, Oyadomari S, Novoa I, Lu P, Zhang Y, Harding HP, et al. Inhibition of a constitutive translation initiation factor 2 alpha phosphatase, CReP, promotes survival of stressed cells. J Cell Biol. 2003;163: 767–775. doi:10.1083/jcb.200308075

19. Connor JH, Weiser DC, Li S, Hallenbeck JM, Shenolikar S. Growth Arrest and DNA Damage-Inducible Protein GADD34 Assembles a Novel Signaling Complex Containing Protein Phosphatase 1 and Inhibitor 1. Mol Cell Biol. 2001;21: 6841–6850. doi:10.1128/MCB.21.20.6841

20. Patterson AD, Hollander MC, Miller GF, Fornace AJ. Gadd34 Requirement for Normal Hemoglobin Synthesis. 2006;26: 1644–1653. doi:10.1128/MCB.26.5.1644

21. Harding HP, Zhang Y, Scheuner D, Chen J, Kaufman RJ, Ron D. Ppp1r15 gene knockout reveals an essential role for translation initiation factor 2 alpha (eIF2a) dephosphorylation in mammalian development. PNAS. 2009;106: 1–6.

22. Grech G, Blázquez-Domingo M, Kolbus A, Bakker WJ, Müllner EW, Beug H, et al. Igbp1 is part of a positive feedback loop in stem cell factor-dependent, selective mRNA translation initiation inhibiting erythroid differentiation. Blood. 2008;112: 2750–60. doi:10.1182/blood-2008-01-133140

23. Horos R, Ijspeert H, Pospisilova D, Sendtner R, Andrieu-Soler C, Taskesen E, et al. Ribosomal deficiencies in Diamond-Blackfan anemia impair translation of transcripts essential for differentiation of murine and human erythroblasts. Blood. 2012;119: 262–72. doi:10.1182/blood-2011-06-358200

24. Ingolia NT, Ghaemmaghami S, Newman JRS, Weissman JS. Genome-wide analysis in vivo of translation with nucleotide resolution using ribosome profiling. Science. 2009;324: 218–23. doi:10.1126/science.1168978

25. Ingolia NT, Lareau LF, Weissman JS. Ribosome profiling of mouse embryonic stem cells reveals the complexity and dynamics of mammalian proteomes. Cell. Elsevier Inc.; 2011;147: 789–802. doi:10.1016/j.cell.2011.10.002

26. Von Lindern M, Deiner EM, Dolznig H, Amelsvoort MP, Hayman MJ, Mullner EW, et al. Leukemic transformation of normal murine erythroid progenitors⊐: v- and c-ErbB act through signaling pathways activated by the EpoR and c-Kit in stress erythropoiesis. Oncogene. 2001;20: 3651–3664.

27. Blazquez-Domingo M, Grech G, Von Lindern M. Translation Initiation Factor 4E Inhibits Differentiation of Erythroid Progenitors. Mol Cell Biol. 2005;25: 8496–506. doi:10.1128/MCB.25.19.8496

28. Pereboom TC, Bondt A, Pallaki P, Klasson TD, Goos YJ, Essers PB, et al. Translation of branched-chain aminotransferase-1 transcripts is impaired in cells haploinsufficient for ribosomal protein genes. Exp Hematol. ISEH − Society for Hematology and Stem Cells; 2014;42: 394–403. doi:10.1016/j.exphem.2013.12.010

29. Salerno F, Paolini NA, Stark R, Von Lindern M, Wolkers MC. Distinct PKC-mediated posttranscriptional events set cytokine production kinetics in CD8+ T cells. PNAS. 2017;114: 9677–9682. doi:10.1073/pnas.1704227114

30. Ingolia NT, Brar G a, Rouskin S, McGeachy AM, Weissman JS. The ribosome profiling strategy for monitoring translation in vivo by deep sequencing of ribosome-protected mRNA fragments. Nat Protoc. 2012;7: 1534–50. doi:10.1038/nprot.2012.086

31. De Klerk E, Fokkema IFAC, Thiadens KAMH, Goeman JJ, Palmblad M, Den Dunnen JT, et al. Assessing the translational landscape of myogenic differentiation by ribosome profiling. Nucleic Acids Res. 2015;43: 4408–4428. doi:10.1093/nar/gkv281

32. Martin M. Cutadapt removes adapter sequences from high-throughput sequencing reads. EMBnet.journal. 2011;17: 10–12. doi:10.14806/ej.17.1.200

33. Dobin A, Davis CA, Schlesinger F, Drenkow J, Zaleski C, Jha S, et al. STAR: Ultrafast universal RNA-seq aligner. Bioinformatics. 2013;29: 15–21. doi:10.1093/bioinformatics/bts635

34. Robinson MD, McCarthy DJ, Smyth GK. edgeR: A Bioconductor package for differential expression analysis of digital gene expression data. Bioinformatics. 2009;26: 139–140. doi:10.1093/bioinformatics/btp616

35. McCarthy DJ, Chen Y, Smyth GK. Differential expression analysis of multifactor RNA-Seq experiments with respect to biological variation. Nucleic Acids Res. 2012;40: 4288–4297. doi: 10.1093/nar/gks042

36. Wildeman M, Van Ophuizen E, Den Dunnen JT, Taschner PEM. Improving Sequence Variant Descriptions in Mutation Databases and Literature Using the Mutalyzer Sequence Variation Nomenclature Checker. Hum Mutat. 2008;29: 6–13. doi:10.1002/humu

37. Bates D, Mächler M, Bolker B, Walker S. Fitting Linear Mixed-Effects Models Using Ime4. J Stat Softw. 2015;67. doi:10.18637/jss.v067.i01

38. Michel AM, Mullan JPA, Velayudhan V, O’Connor PBF, Donohue CA, Baranov P V. RiboGalaxy: A browser based platform for the alignment, analysis and visualization of ribosome profiling data. RNA Biol. Taylor & Francis; 2016;13: 316–319. doi: 10.1080/15476286.2016.1141862

39. Lareau LF, Hite DH, Hogan GJ, Brown PO. Distinct stages of the translation elongation cycle revealed by sequencing ribosome-protected mRNA fragments. Elife. 2014;2014: 1–16. doi:10.7554/eLife.01257

40. Gerashchenko M V., Gladyshev VN. Translation inhibitors cause abnormalities in ribosome profiling experiments. Nucleic Acids Res. 2014;42: e134. doi:10.1093/nar/gku671

41. Simsek D, Tiu GC, Flynn RA, Xu AF, Chang HY, Barna M, et al. The Mammalian Ribo-interactome Reveals Ribosome Functional Diversity and Heterogeneity. Cell. Elsevier Inc.; 2017;169: 1051–1057.e18. doi:10.1016/j.cell.2017.05.022

42. Prostko CR, Brostrom MA, Brostrom CO. Reversible phosphorylation of eukaryotic initiation factor 2 alpha in response to endoplasmic reticular signaling. Mol Cell Biochem. 1993;127–128: 255-65. Available: http://www.ncbi.nlm.nih.gov/pubmed/7935356

43. Hetz C. The unfolded protein response: controlling cell fate decisions under ER stress and beyond. Nat Rev Mol Cell Biol. Nature Publishing Group; 2012;13: 89–102. doi: 10.1038/nrm3270

44. Stöckel D, Kehl T, Trampert P, Schneider L, Backes C, Ludwig N, et al. Multiomics enrichment analysis using the GeneTrail2 web service. Bioinformatics. 2016;32: 1502–1508.

45. Malhotra JD, Kaufman RJ. ER Stress and Its Functional Link to Mitochondria⊐: Role in Cell Survival and Death. Cold Spring Harb Perspect Biol. 2011;3: a004424.

46. Fujishima N, Hirokawa M, Aiba N, Ichikawa Y, Fujishima M, Komatsuda A, et al. Gene Expression Profiling of Human Erythroid Progenitors by Micro-Serial analysis of Gene Expression. Int J Hematol. 2004;80: 239–45.

47. Michel AM, Fox G, Kiran AM, BoC De, Connor PBFO, Heaphy SM, et al. GWIPS-viz⊐: development of a ribo-seq genome browser. Nucleic Acids Res. 2014;42: 859–864. doi: 10.1093/nar/gkt1035

48. Cho H, Kim KM, Kim YK. Human Proline-Rich Nuclear Receptor Coregulatory Protein 2 Mediates an Interaction between mRNA Surveillance Machinery and Decapping Complex. Mol Cell. Elsevier Inc.; 2009;33: 75–86. doi:10.1016/j.molcel.2008.11.022

49. Baird TD, Palam LR, Fusakio ME, Willy JA, Davis CM, McClintick JN, et al. Selective mRNA translation during eIF2 phosphorylation induces expression of IBTKα. Mol Biol Cell. 2014;25: 1686–97. doi: 10.1091/mbc.E14-02-0704

50. Hinnebusch AG. Translational regulation of yeast GCN4. J Biol Chem. 1997;272: 21661–21664. Available: http://www.jbc.org/content/272/35/21661.short

51. Nanda JS, Saini AK, Muñoz AM, Hinnebusch AG, Lorsch JR. Coordinated movements of eukaryotic translation initiation Factors eIF1, eIF1A, and eIF5 trigger phosphate release from eIF2 in response to start codon recognition by the ribosomal preinitiation complex. J Biol Chem. 2013;288: 5316–5329. doi:10.1074/jbc.M112.440693

52. Sendoel A, Dunn JG, Rodriguez EH, Naik S, Gomez NC, Hurwitz B, et al. Translation from unconventional 5’ start sites drives tumour initiation. Nature. Nature Publishing Group; 2017;541: 494–499. doi:10.1038/nature21036

53. Golovko A, Kojukhov A, Guan BJ, Morpurgo B, Merrick WC, Mazumder B, et al. The eIF2A knockout mouse. Cell Cycle. Taylor & Francis; 2016;15: 3115–3120. doi:10.1080/15384101.2016.1237324

54. Reid DW, Chen Q, Tay AS, Shenolikar S, Nicchitta C V. The Unfolded Protein Response Triggers Selective mRNA Release from the Endoplasmic Reticulum. Cell. Elsevier Inc.; 2014;158: 1362–1374. doi:10.1016/j.cell.2014.08.012

55. Reid DW, Tay ASL, Sundaram JR, Lee ICJ, Chen Q, George SE, et al. Complementary Roles of GADD34- and CReP-Containing Eukaryotic Initiation Factor 2 α Phosphatases during the Unfolded Protein Response. Mol Cell Biol. 2016;36: 1868–1880. doi:10.1128/MCB.00190-16.Address

56. Fritsch C, Herrmann A, Nothnagel M, Szafranski K, Huse K, Schumann F, et al. Genome-wide search for novel human uORFs and N-terminal protein extensions using ribosomal footprinting. Genome Res. 2012;22: 2208–2218. doi:10.1101/gr.139568.112.2208

57. Cho H, Park OH, Park J, Ryu I, Kim J, Ko J, et al. Glucocorticoid receptor interacts with PNRC2 in a ligand-dependent manner to recruit UPF1 for rapid mRNA degradation. PNAS. 2015;112: 1540–1549. doi:10.1073/pnas.1409612112

58. Bauer A, Tronche F, Wessely O, Kellendonk C, Reichardt HM, Steinlein P, et al. The glucocorticoid receptor is required for stress erythropoiesis. Genes Dev. 1999;13: 2996–3002.

59. Jacobs-Helber SM, Abutin RM, Tian C, Bondurant M, Wickrema A, Sawyer ST. Role of JunB in erythroid differentiation. J Biol Chem. 2002;277: 4859–4866. doi:10.1074/jbc.M107243200

60. Gurzov E, Ortis F, Cunha D, Gosset G, Li M, Cardozo A, et al. Signaling by IL-1 b + IFN-⊐ and ER stress converge on DP5 / Hrk activation⊐: a novel mechanism for pancreatic β-cell apoptosis. Cell Death Differ. 2009;16: 1539–1550. doi:10.1038/cdd.2009.99

61. Dragon S, Offenhauser N, Baumann R. Fos Expression in Erythroid Cells of the Chick Embryo. Am J Physiol Regul Integr Comp Physiol. 2002;282: R1219–26.

62. Sebastian S, Sreenivas P, Sambasivan R, Cheedipudi S, Kandalla P, Pavlath GK, et al. MLL5, a trithorax homolog, indirectly regulates H3K4 methylation, represses cyclin A2 expression, and promotes myogenic differentiation. Proc Natl Acad Sci. 2009;106: 4719–4724. doi:10.1073/pnas.0807136106

